# *Mycobacterium tuberculosis* Modulates the Metabolism of Alternatively Activated Macrophages to Promote Foam Cell Formation and Intracellular Survival

**DOI:** 10.1101/2019.12.13.876300

**Authors:** Melanie Genoula, José Luis Marín Franco, Mariano Maio, Belén Dolotowicz, Malena Ferreyra, M. Ayelén Milillo, Rémi Mascarau, Eduardo José Moraña, Domingo Palmero, Federico Fuentes, Beatriz López, Paula Barrionuevo, Olivier Neyrolles, Céline Cougoule, Geanncarlo Lugo-Villarino, Christel Vérollet, María del Carmen Sasiain, Luciana Balboa

## Abstract

The ability of Mycobacterium tuberculosis (Mtb) to persist inside host cells relies on metabolic adaptation, like the accumulation of lipid bodies (LBs) in the so-called foamy macrophages (FM). Indeed, FM are favorable to Mtb. The activation state of macrophages is tightly associated to different metabolic pathways, such as lipid metabolism, but whether differentiation towards FM differs between the macrophage activation profiles remains unclear. Here, we aimed to elucidate if distinct macrophage activation states exposed to a tuberculosis-associated microenvironment can accumulate LBs, and its impact on the control of infection. We showed that signal transducer and activator of transcription 6 (STAT6) activation in interleukin (IL)-4-activated human macrophages (M(IL-4)) prevents FM formation induced by pleural effusion from patients with tuberculosis. In these cells, LBs are disrupted by lipolysis, and the released fatty acids enter the β-oxidation (FAO) pathway fueling the generation of ATP in mitochondria. We demonstrated that inhibition of the lipolytic activity or of the FAO drives M(IL-4) macrophages into FM. Also, exhibiting a predominant FAO metabolism, mouse alveolar macrophages are less prone to become FM compared to bone marrow derived-macrophages. Upon Mtb infection, M(IL-4) macrophages are metabolically re-programmed towards the aerobic glycolytic pathway and evolve towards a foamy phenotype, which could be prevented by FAO activation or inhibition of the hypoxia-inducible factor 1-alpha (HIF-1α)-induced glycolytic pathway. In conclusion, our results demonstrate a role for STAT6-driven FAO in preventing FM differentiation, and reveal an extraordinary capacity by Mtb to rewire metabolic pathways in human macrophages and induce the favorable FM.

**IMPORTANCE:** Tuberculosis (TB) is an infectious disease caused by Mycobacterium tuberculosis (Mtb). While its treatment was already standardized, TB remains one of the top 10 death causes worldwide. A major problem is the efficient adaptation of Mtb to the macrophage intracellular milieu, which includes deregulation of the lipid metabolism leading to the formation of foamy macrophages (FM) which are favorable to Mtb. A critical aspect of our work is the use of tuberculous pleural effusions (TB-PE) — human-derived biological fluid capable of mimicking the complex microenvironment of the lung cavity upon Mtb infection — to study the FM metabolic modulation. We revealed how the STAT6 transcription factor prevents FM formation induced by PE-TB, and how Mtb counteracts it by activating another transcription factor, HIF-1α, to re-establish FM. This study provides key insights in host lipid metabolism, macrophage biology and pathogen subversion strategies, to be exploited for prevention and therapeutic purposes in infectious diseases.

## INTRODUCTION

Tuberculosis (TB) is a highly contagious disease caused by Mycobacterium tuberculosis (Mtb). Even though the treatment of the disease has been standardized for a while, TB still remains one of the top 10 causes of death worldwide (1). Chronic host-pathogen interaction in TB leads to extensive metabolic remodeling in both the host and the pathogen (2). The success of Mtb as a pathogen is in a large part due to its efficient adaptation to the intracellular milieu of human macrophages, leading to metabolic changes in infected host cells. One of these changes is the dysregulation of the lipid metabolism, which induces the formation of foamy macrophages (FM). FM are cells filled of lipid bodies (LBs) that are abundant in granulomatous structures in both experimentally infected animals and patients (3, 4) and fail to control the infection (5). Recently, we used the pleural effusions (PE) from TB patients (TB-PE) as a tool to recapitulate human lung TB-associated microenvironment and demonstrated that uninfected-macrophages exposed to TB-PE form FM displaying an immunosuppressive profile though the activation of the interleukin (IL)-10/STAT3 axis (6).

It is widely accepted that macrophages undergo different activation programs by which they carry out unique physiological and defensive functions. Essentially, macrophages can modify their metabolic functions from a healing/repairing/growth promoting setting (M2 macrophages) towards a killing/inhibitory/microbicidal profile (M1 macrophages) (7, 8), representing opposing ends of the full activation spectrum. The M1 macrophages, generally induced by interferon (IFN)-γ and/or lipopolysaccharide (LPS) stimulation, are endowed with microbicidal properties; the M2 macrophages, usually differentiated upon IL-4 or IL-13 stimulation, are reported to be immunomodulatory and poorly microbicidal, resulting in impaired anti-mycobacterial properties (9–11). In addition, while M1 macrophages rely on glycolysis dependent on the hypoxia-inducible factor 1-alpha (HIF-1α) activation (12), M2 macrophages require the induction of fatty acid oxidation (FAO), at least in the murine model, (13–16) through signal transducer and activator of transcription 6 (STAT6) activation (17). FAO is the mitochondrial process of breaking down a fatty acid into acetyl-CoA units, and requires the carnitine palmitoyltransferase (CPT) system, which consists of one transporter and two mitochondrial membrane enzymes: CPT1 and CPT2 (18). This facilitates the transport of long-chain fatty acids into the mitochondrial matrix, where they can be metabolized by the oxidative phosphorylation (OXPHOS) pathway and produce ATP. Moreover, in the context of Mtb infection, macrophages have been recently shown to be pre-programmed towards different metabolic pathways, according to their ontogeny. Indeed, recruited lung interstitial macrophages are glycolytically active and capable of controlling Mtb infection, while resident alveolar macrophages are committed to FAO and OXPHOS bearing higher bacillary loads (19).

Considering that mitochondria are the main site of lipid degradation, and that mitochondrial metabolic functions are known to differ between macrophages profiles, we investigated if the different activation programs of human macrophages differ in their ability to form FM, and whether this impacts the control of Mtb growth. To this end, we employed our already characterized model of TB-induced FM differentiation, using TB-PE (6), and explored the molecular and metabolic pathways capable of altering the accumulation of LBs. The pleural effusion is an excess of fluid recovered from pleural space characterized by a high protein content and specific leukocytes (20). We have previously demonstrated that the TB-PE tilted ex vivo human monocyte differentiation towards an anti-inflammatory M2-like macrophage activation, reproducing the phenotype exhibited by macrophages directly isolated from the pleural cavity of TB patients and from lung biopsies of non-human primates with advance TB (21). Likewise, we showed that the acellular fraction of TB-PE modified the lipid metabolism of human macrophages, leading them into FM and impacting their effector functions against the bacilli (6). In this work, we demonstrated that STAT6-driven FAO prevents FM differentiation in macrophages exposed to the TB-PE and, conversely, we revealed how Mtb counteracts it by inhibiting another transcription factor, HIF-1α, to re-establish FM. Therefore, this study contributes to our understanding of how alterations of the host metabolic pathways affect pathogen persistence.

## RESULTS

### STAT6 activation in M(IL-4) macrophages prevents the formation of TB-induced foamy macrophages

In order to evaluate whether different activation programs in human macrophages differ in their ability to form FM upon treatment with the acellular fraction of tuberculous pleural effusions (TB-PE), macrophages were differentiated with IL-4, IL-10 or IFN-γ, and treated (or not) with TB-PE. The intracellular accumulation of LBs was then determined by Oil Red O (ORO) staining. The profiles of macrophages activation were evaluated by determining the expression of cell-surface and other markers (Fig. S1A-B). As expected, PE-TB induced FM formation in non-polarized (M0) macrophages (6), (Fig. 1A). Interestingly, TB-PE also induced the accumulation of LBs in M(IFN-γ) and M(IL-10), but not in M(IL-4) macrophages, suggesting that the activation program can influence PE-TB-induced LB formation. In agreement with our previous work (6), the accumulation of LBs observed in M0, M(IFN-γ) and M(IL-10) cells was specific for TB-PE treatment since PE from patients with heart-failure (HF-PE) did not show LBs formation (Fig. 1A). Moreover, the addition of cytokines known to drive the alternative activation program in macrophages, i.e. recombinant IL-4 or IL-13 (Fig. S2A), inhibited the accumulation of LBs in TB-PE-treated M0 in a dose-dependent manner (Fig. 1B-C and S2B-C). Since IL-4/IL-13-dependent activation of STAT6 is required for alternative macrophage activation (22, 23), we evaluated the impact of STAT6 inhibition in M(IL-4) macrophages on LB accumulation by using the chemical inhibitor AS1517499, which prevents STAT6 phosphorylation (Fig. S2D-E and (24)). Inhibition of STAT6 activation enabled M(IL-4) cells to accumulate LBs (Fig. 1D).

**Figure 1.**
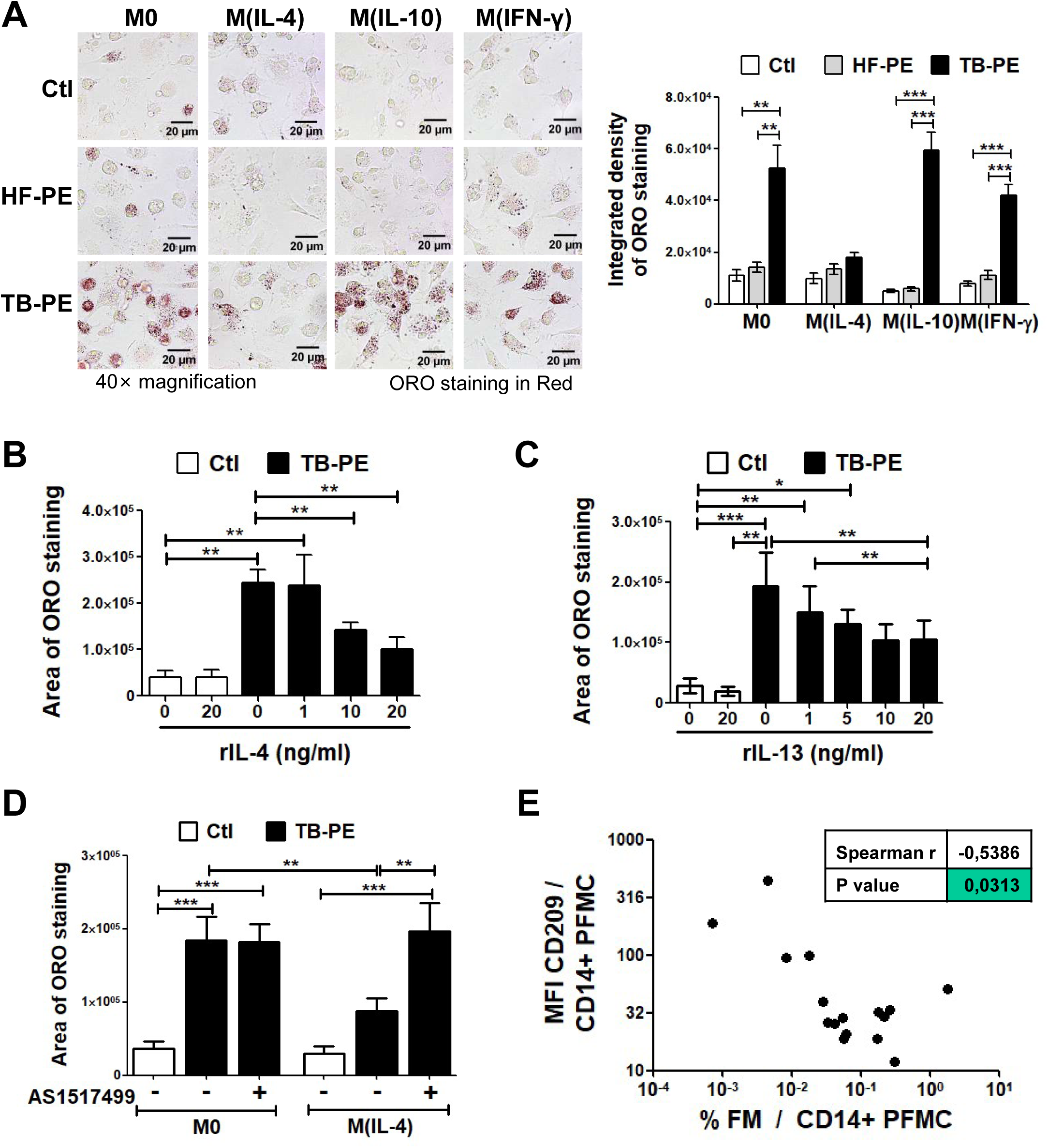
The IL-4/STAT6 Axis prevents the formation of foamy macrophages. Human macrophages were left untreated (M0) or polarized either IL-4 (M(IL-4)), IL-10 (M(IL-10)) or IFN-γ (M(IFN-γ)) for 48 h, treated or not with the acellular fraction of TB pleural effusions (TB-PE) or heart-failure-associated effusions (HF-PE) for 24 h and then stained with Oil Red O (ORO). **(A)** Left panel: Representative images (40× magnification), right panel: the integrated density of ORO staining. (B-C) Quantification of area of ORO staining of macrophages polarized with either different doses of recombinant IL-4 **(B)** or IL-13 **(C)** for 48 h and exposed to TB-PE for further 24 h. **(D)** Quantification of area of ORO staining of M0 and M(IL-4) macrophages treated with TB-PE and exposed or not to AS1517499, a chemical inhibitor of STAT6. Values are expressed as means ± SEM of six independent experiments, considering five microphotographs per experiment. Friedman test followed by Dunn’s Multiple Comparison Test: *p<0.05; **p<0.01; ***p<0.001 as depicted by lines. **(E)** Correlation study between the mean fluorescence intensity (MFI) of CD209 cell-surface expression in CD14^+^ cells from TB pleural cavity and the percentage of lipid-laden CD14^+^ cells within the pleural fluids mononuclear cells (PFMC) (n=16) found in individual preparations of TB-PE. Spearman’s rank test.

In order to evaluate whether M(IL-4) macrophages were less prone to become foamy in vivo during TB, we assessed the expression levels of surface markers in CD14^+^ cells as well as the percentages of FM found within the pleural fluid mononuclear cells isolated from TB patients. We found that the expression levels of an IL-4-driven marker signature, such as CD209/DC-SIGN, was negatively correlated with the percentage of FM found among the pleural fluid mononuclear cells, unlike other surface receptors (Fig. 1E and S3). Noticeably, we observed a positive correlation between the percentage of FM and the expression of MerTK, a M(IL-10)-associated marker(Fig. S3), which confirms our previous results showing that IL-10 enhances FM formation in TB-PE-treated macrophages (6). These results indicate that the IL-4-driven phenotype prevents the foamy program in human macrophages in the context of a natural Mtb infection. Therefore, the STAT6 signaling pathway mediates the inhibition of FM formation induced by TB-PE.

### The biogenesis of LBs is not impaired in TB-PE-treated M(IL-4) macrophages

In order to elucidate how the IL-4/STAT6 axis could interfere with PE-TB-induced macrophage differentiation into FM, we investigated cellular mechanisms associated with the biogenesis of LBs. We found that M0 and M(IL-4) macrophages did not differ in their ability to uptake fatty acids (Fig. 2A). In addition, TB-PE was able to induce the expression of the scavenger receptor CD36, which binds long-chain fatty acids and facilitates their transport into cells (Fig. 2B), and that of Acyl-CoA acetyltransferase (ACAT), the enzyme that esterifies free cholesterol (Fig. 2C). We next evaluated the morphometric features of the LBs formed in TB-PE-stimulated M0 and M(IL-4) cells. As shown in figure 2D, although LBs are formed within M(IL-4) cells, they were significantly smaller. We also confirmed the presence of smaller LBs in TB-PE-treated M(IL-4) macrophages compared to M0 cells by transmission electron microscopy (Fig. 2E, arrows). These results suggest that while the biogenesis of LBs is not impaired, they are smaller in TB-PE-exposed M(IL-4) macrophages compared to TB-PE-treated M0 macrophages.

**Figure 2.**
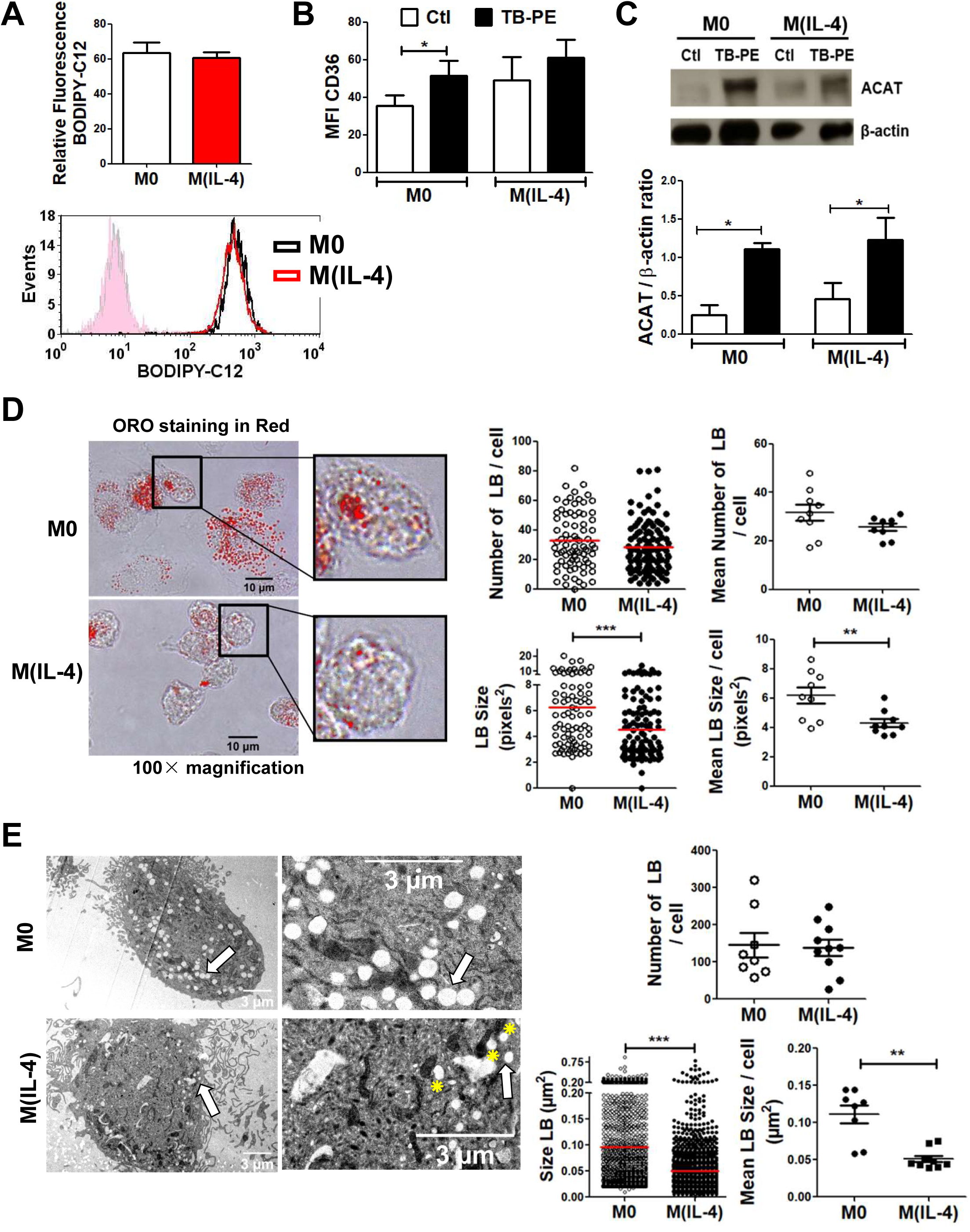
The biogenesis of LBs is not impaired in TB-PE-treated M(IL-4) macrophages. **(A)** Human macrophages were left untreated (M0) or polarized with IL-4 (M(IL-4)) for 48 h and then exposed or not to the red-labelled saturated fatty acid C12 (BODIPY C12) for 1 min. Upper panel: relative BODIPY C12 fluorescence, lower panel: representative histogram. Values are expressed as means ± SEM of four independent experiments. **(B)** Median fluorescence intensity (MFI) of CD36 measured by flow cytometry in M0 and M(IL-4) macrophages exposed or not to TB-PE. Values are expressed as means±SEM of six independent experiments. Friedman test followed by Dunn’s Multiple Comparison Test: *p<0.05 for experimental condition vs Ctl. **(C)** Upper panel: analysis of ACAT and β-actin protein level by Western Blot; lower panel: quantification in M0 and M(IL-4) macrophages treated or not with TB-PE for 24 h (n=4). Wilcoxon signed rank test: *p<0.05 as depicted by lines. **(D)** Morphometric analysis of LBs in ORO-labelled TB-PE-treated M0 and M(IL-4) macrophages. Representative images of ORO staining (100× magnification) are shown. Number of LB (upper left panel) and size (lower left panel) per cell are shown, and their mean is represented by red bars. Mann Whitney test: ***p<0.001. Mean number of LB (upper right panel) and mean size (lower right panel) per cell per donor (n=9 donors). Each determination represents the mean of 20-40 individual cells per donor. Wilcoxon signed rank test: **p<0.01. **(E)** Electron microscopy micrographs of TB-PE-treated M0 and M(IL-4) macrophages showing LBs (white arrows) and LBs nearby mitochondria (yellow asterisks). Upper panel: numbers of LB per cell; Lower panels: size of LB in TB-PE-treated M0 and M(IL-4) macrophages considering all LB’ size determinations (left) or the mean of 8-10 individual cells of one representative donor (right). Mann Whitney test: **p<0.01; ***p<0.001.

### Enhanced lipolysis mediates the inhibition of the accumulation of LBs in M(IL-4) macrophages

As lysosomal lipolysis is essential for the alternative activation of macrophages (14), we wondered whether lipolysis could be a mechanism by which IL-4/IL-13 reduce LB accumulation induced by TB-PE. We found that both M(IL-4) and M(IL-13) cells displayed higher lipolytic activity than M0 cells, bearing the peak of lipolytic activity at 24 h post-IL-4 addition, as measured by glycerol release (Fig. 3A **left panel and S4A**). Particularly, TB-PE-treated M(IL-4) cells released higher levels of glycerol than untreated M(IL-4) cells or TB-PE-treated M0 macrophages, indicating that more triglycerides were broken down in M(IL-4) macrophages upon TB-PE treatment (Fig. 3A **right panel**). Pharmacological inhibition of STAT6 activity reduced the release of glycerol in TB-PE-treated M(IL-4) cells (Fig. 3B). Thereafter, we found that inhibition of lipase activity by the drugs Orlistat, which targets neutral lipolysis, or Lalistat, which targets acid lipolysis, enabled M(IL-4) macrophages to accumulate LBs and reduce glycerol release (Fig. 3C-D **and S4B-C**). Hence, the activation of the IL4/STAT6 axis in M(IL-4) macrophages promotes lipolysis impairing the accumulation of TB-PE-induced LBs.

**Figure. 3.**
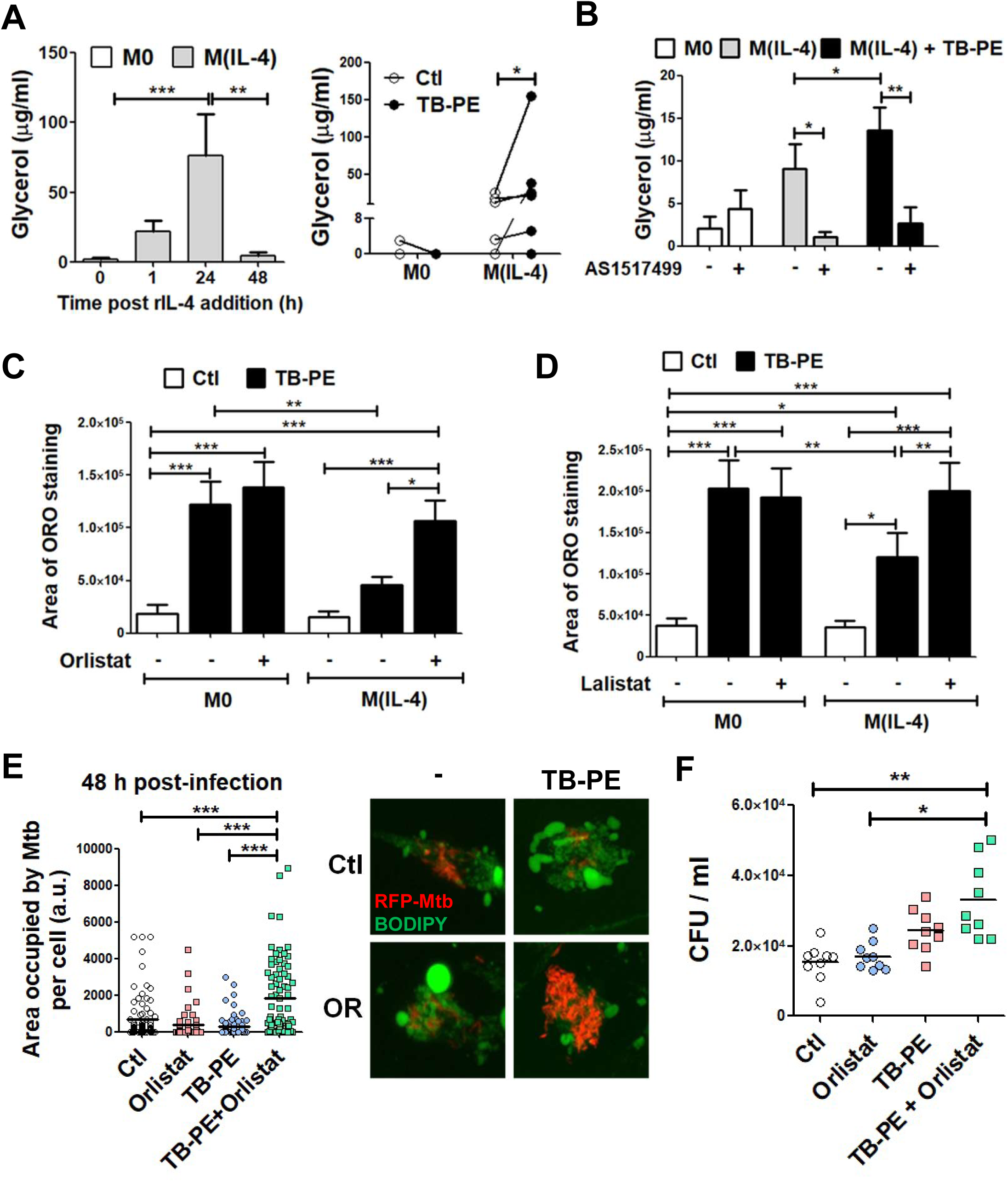
Enhanced lipolysis inhibits lipid bodies’ accumulation in M(IL-4) macrophages. **(A)** Macrophages were polarized or not with IL-4 for 1, 24 or 48 h (left panel), and then treated or not with TB-PE for further 24 h (right panel). Thereafter, culture-medium was replaced by PBS containing 2% BSA (fatty acid-free), and supernatants were collected 6h later to measure the release of glycerol by colorimetric assays. Left panel: kinetics of glycerol release after IL-4 addition (n=10). Right panel: glycerol release after TB-PE treatment in M0 or M(IL-4) macrophages (n=6). **(B)** Glycerol release by macrophages exposed or not to AS1517499 (n=7). Values are expressed as means ± SEM of independent experiments. Friedman test followed by Dunn’s Multiple Comparison Test: *p<0.05 as depicted by lines. (C-D) ORO staining of M0 and M(IL-4) macrophages treated with TB-PE and exposed or not to either Orlistat **(C)** or Lalistat (D). Values are expressed as means ± SEM of five **(C)** or seven **(D)** independent experiments, considering five microphotographs per experiment. Friedman test followed by Dunn’s Multiple Comparison Test: *p<0.05; **p<0.01; ***p<0.001 as depicted by lines. (E-F) M(IL-4) macrophages were treated or not with TB-PE in the presence or not of Orlistat for 24 h, washed and infected with RFP-Mtb **(E)** or unlabeled Mtb (F). **(E)** Area with RFP-Mtb per cell in z-stacks from confocal laser scanning microscopy images at 48 h post-infection. Representative images are shown (60× magnification). Each determination represents individual cells of one donor. One way-ANOVA followed by Bonferroni test: ***p<0.001. **(F)** Intracellular colony forming units determined at 48 h post-infection (n=9). Friedman followed by Dunn’s Multiple Comparison Test: *p<0.05; **p<0.01 as depicted by lines.

Next, we wondered whether the differentiation of M(IL-4) macrophages into FM could impact on the control of the bacillary loads. To this end, we determined the bacterial content in TB-PE-treated M(IL-4) macrophages, which have been exposed (or not) to Orlistat prior to Mtb infection, with two experimental approaches by: i/ assessing the area of red fluorescent protein (RFP)-Mtb associated to individual cells by confocal microscopy, or ii/ enumerating the CFU recovered from the cultures. We confirmed that the M(IL-4) cells treated with TB-PE in the presence of Orlistat displayed LBs after 2 h post-infection (Fig S4D). While the entrance of mycobacteria at early time points was comparable among conditions **(Fig S4D-E)**, treatment of M(IL-4) cells with TB-PE in the presence of Orlistat rendered the cells more susceptible to intracellular Mtb replication (Fig. 3E-F **and S4E**). Therefore, inhibition of lipolysis prior to infection leads M(IL-4) cells to become foamy, and is associated with an increase in Mtb intracellular growth.

### The inhibition of fatty acid transport into mitochondria allows lipid accumulation within LBs in TB-PE-treated M(IL-4) macrophages

Aside from storage in lipid droplets, fatty acids can also be transported into the mitochondria and be oxidized into acetylCoA by FAO. In this regard, the ultrastructural analysis of the cells revealed that LBs were frequently found nearby mitochondria in TB-PE-treated M(IL-4) macrophages (Fig. 2E, asterisks). Therefore, we hypothesized that the enhanced lipolytic activity in M(IL-4) cells might drive fatty acids into the mitochondria to be used for FAO leading to high ATP production through the OXPHOS pathway. The expression of CPT1, an enzyme which is considered to be rate limiting for β-oxidation of long-chain fatty acids (18), was evaluated. As expected, higher levels of CPT1A mRNA in M(IL-4) compared to M0 macrophages were observed. Although the presence of TB-PE diminishes CPT1A expression, TB-PE-treated M(IL-4) cells still conserve higher levels than TB-PE-treated M0 cells (Fig. 4A). Moreover, we performed a pulse-chase assay to track fluorescent fatty acids in relation to LBs and mitochondria in TB-PE-treated M0 and M(IL-4) macrophages. We found that the saturated fatty acid Red C12 was more often accumulated in neutral lipids within LBs in TB-PE-treated M0 cells, while more Red C12 signal was accumulated into mitochondria in TB-PE-treated M(IL-4) macrophages (Fig. 4B).

**Figure 4.**
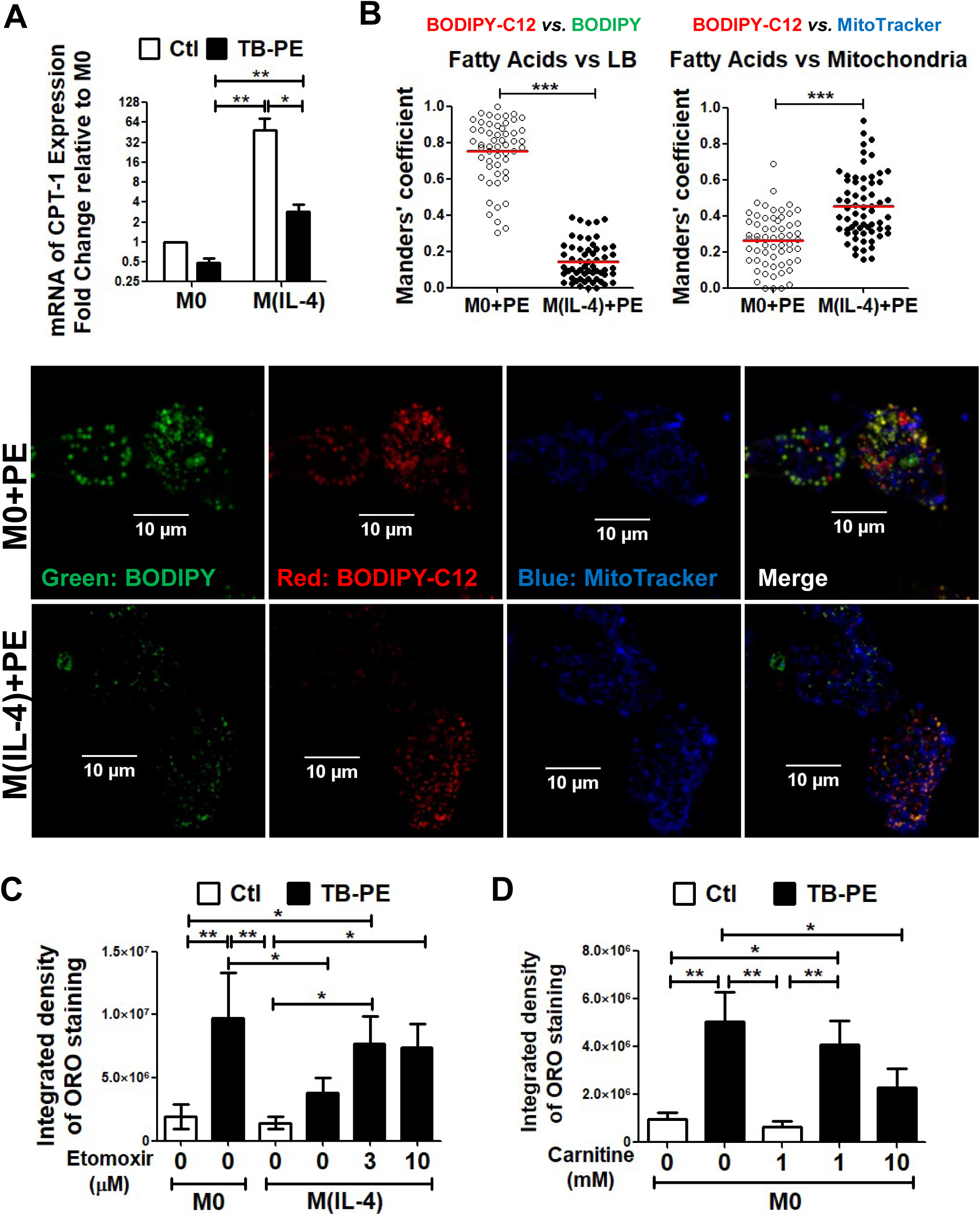
Inhibition of fatty acid transport into mitochondria allows lipids accumulation within LBs in TB-PE-treated M(IL-4) macrophages. M0 and M(IL-4) macrophages treated or not with TB-PE for 24 h **(A)** mRNA expression of CPT1A relative to M0 (n=8). **(B)** Cells were pulsed with BODIPY Red C12 overnight during TB-PE treatment, then LBs were labeled using Green-BODIPY 493/503, and mitochondria were labeled using Far Red-MitoTracker. Quantification of the co-occurrence fractions of fatty acids (C12) with either LBs or Mitochondria in TB-PE-treated M0 and M(IL-4) macrophages were quantified by Manders’ coefficient analysis. Mann Whitney test: ***p<0.001. Representative images of single and merged channels are shown. **(C)** ORO staining of M0 and M(IL-4) macrophages treated with TB-PE and exposed or not to Etomoxir. Values are expressed as means ± SEM of six independent experiments, considering five microphotographs per experiment. **(D)** ORO staining of M0 macrophages treated with TB-PE and exposed or not to L-carnitine. Values are expressed as means ± SEM of five independent experiments, considering five microphotographs per experiment. Friedman test followed by Dunn’s Multiple Comparison Test: *p<0.05; **p<0.01 for experimental condition vs Ctl or as depicted by lines.

We next decided to pharmacologically inhibit the activity of CPT1 and evaluate its impact on FM formation. We used an inhibitor of FAO (Etomoxir) in M(IL-4) cells at doses (3-10 μM) that do not induce cell death (Fig. S5A) or off-target effects (25). We found that this treatment could drive TB-PE-treated M(IL-4) cells to accumulate LBs (Fig. 4C). In line with this, when we foment the FAO pathway by adding exogenous L-carnitine, which conjugates to fatty acids allowing them to enter mitochondria (26), FM formation was inhibited in TB-PE-treated M0 cells (Fig. 4D **and S5B**). Altogether, these results indicate that IL-4 enhances fatty acid transport into the mitochondria thus reducing lipid accumulation within LB structures in TB-PE-treated M(IL-4) macrophages.

### An oxidative metabolism in macrophages is associated with less accumulation of LBs

Since M(IL-4) macrophages oxidize fatty acids to fuel the OXPHOS pathway, we hypothesized that their refractoriness to accumulate fatty acids and become FM could be associated to their particular IL-4-driven metabolism. First, we demonstrated that the M(IL-4) phenotype was still conserved upon TB-PE treatment as judged by the expression of CD209, CD200R, and pSTAT6 (Fig. S5C-D). Next, we evaluated the effect of TB-PE treatment on a key function associated to the M(IL-4) profile, such as the uptake of apoptotic polymorphonuclear leukocytes, i.e. efferocytosis. As observed in **figure S5E**, TB-PE-treated M(IL-4) cells were even more efficient at efferocytosis than untreated-M(IL-4) cells. As IL-4 levels in TB-PE were undetectable by ELISA (less than 6 pg/ml, data not shown), the observed effect could not be due to IL-4 within TB-PE. In line with this, M(IL-10) or M(IFN-γ) macrophages did not show detectable expression of the phosphorylated form of STAT6 in the presence of TB-PE (Fig. S5D). Hence, important features associated with the alternative activation driven by IL-4 were not impaired by TB-PE treatment. Moreover, mitochondrial respiration is associated with M(IL-4) macrophages (22). In this regard, we found a higher mitochondrial respiration in TB-PE-treated M(IL-4) macrophages in comparison to untreated cells (Fig. 5A-B). We also measured the release of lactate as evidence of the activation of the glycolytic pathway, and found that lactate release was lower in M(IL-4) cells regardless its treatment with TB-PE compared to M(IFN-γ) cells, which are known to be glycolytic cells (Fig. 5C and (27)). The glycolytic pathway is highly regulated by HIF-1α activity (12, 28). In fact, the stabilization of HIF-1α expression resulted in both the upregulation of glycolysis and the suppression of FAO (29). For this reason, we next treated M0 and M(IL-4) with dimethyloxaloylglycine (DMOG), which leads to HIF-1α stabilization (30), and evaluated its impact on FM formation upon TB-PE treatment. As shown in figure 5D and S6A-B, an increase in LBs accumulation was observed after HIF-1α stabilization. Importantly, when we looked at murine alveolar macrophages (AM), which are known to be committed to an FAO and OXPHOS pathway (19), we observed that they did not accumulate LBs in response to mycobacterial lipids in comparison to bone marrow derived macrophages (BMDM) (Fig. 5E). By contrast, when exposed to either a HIF-1α potentiator (DMOG) or a FAO inhibitor (Etomoxir), AM did become foamy (Fig. 5F and S6C). Of note, murine IL-4-activated BMDM were less prone to become foamy upon mycobacterial lipids exposure compared to M0 macrophages (Fig. 5G and S6D). These results demonstrate that an oxidative metabolic state in alternatively activated macrophages is associated with less LBs accumulation and FM formation.

**Figure 5.**
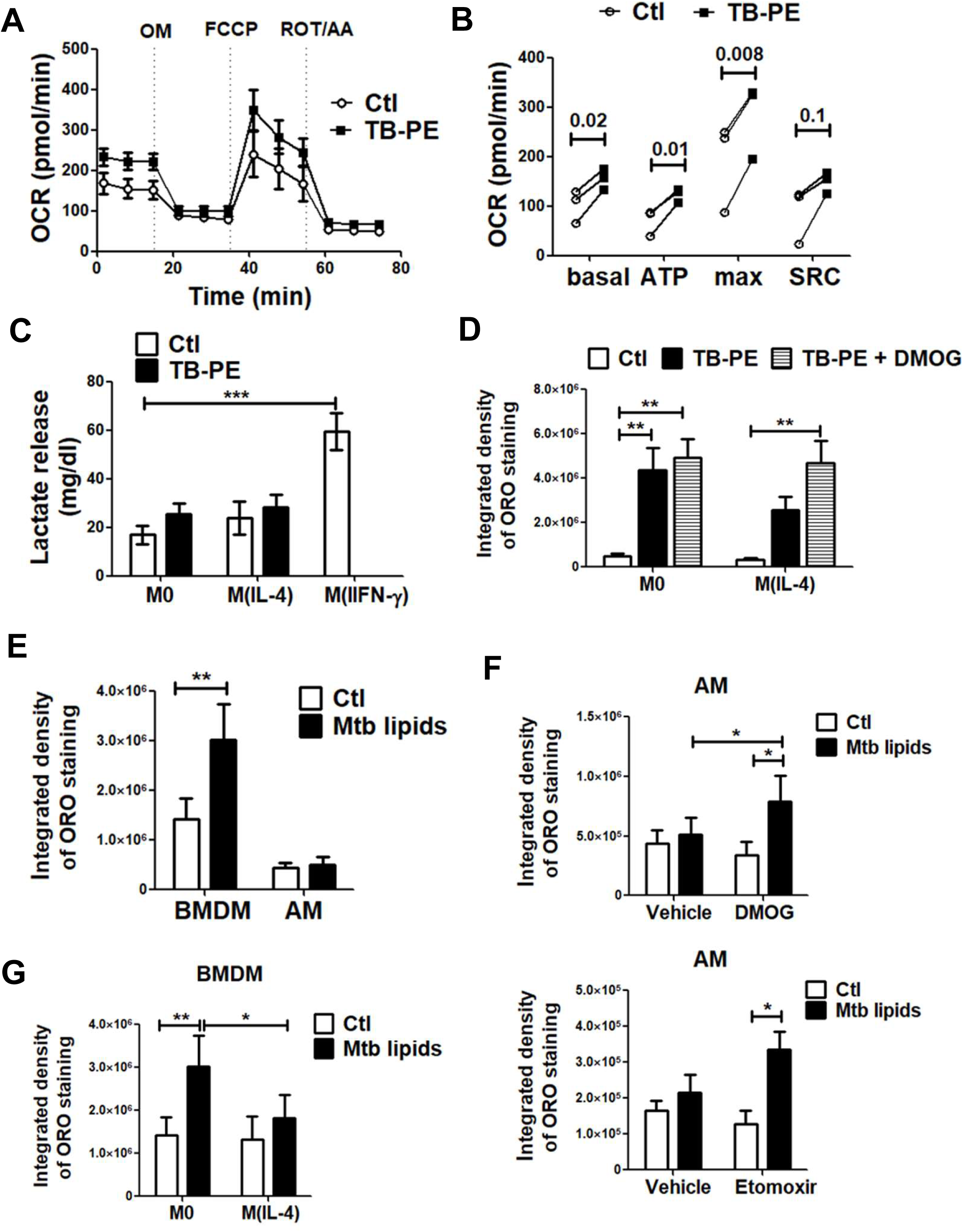
An oxidative metabolism is associated with less accumulation of lipid bodies. **(A-B)** Mitochondrial respiration in M(IL-4) cells exposed or not to TB-PE for 24 h. **(A)** Changes in oxygen consumption rate (OCR) in response to sequential injections of oligomycin (OM), carbonyl cyanide 4-(trifluoromethoxy)phenylhydrazone (FCCP), and rotenone (ROT) + antimycin A (AA) were measured by using an extracellular flux analyzer. Result showed was one representative from three independent experiments. **(B)** Calculated basal respiration, ATP production, maximal respiration and spare respiratory capacity are plotted in bar graphs. Values are expressed as means of triplicates and three independent experiments are shown. T-tests were applied. **(C)** Lactate release by M(IL-4) cells treated or not with TB-PE and by M(IFN-γ) as positive control. Values are expressed as means ± SEM of nine independent experiments. Friedman test followed by Dunn’s Multiple Comparison Test: ***p<0.001 for M(IFN-γ) vs Ctl. **(D)** Integrated density of ORO staining of M0 and M(IL-4) macrophages treated with TB-PE in the presence or not of DMOG. Values are expressed as means ± SEM of 6 independent experiments, considering five microphotographs per experiment. Friedman test followed by Dunn’s Multiple Comparison Test: *p<0.05; for experimental condition vs Ctl or as depicted by lines. **(E)** Integrated density of ORO staining of murine bone marrow derived macrophages (BMDM) and alveolar macrophages (AM) treated with a total lipids’ preparation from Mtb (Mtb lipids). **(F)** Integrated density of ORO staining of murine AM treated or not with Mtb lipids in the presence of DMOG (upper panel) or Etomoxir (lower panel). Values are expressed as means ± SEM of four independent experiments, considering five microphotographs per experiment. One-way ANOVA followed by Bonferroni post-test: *p<0.05; **p<0.01 as depicted by lines. **(G)** Murine bone marrow derived macrophages (BMDM) were left untreated (M0) or polarized with IL-4 (M(IL-4)) for 48 h, treated with Mtb lipids for further 24 h, and ORO-stained for quantification of LBs accumulation. Values are expressed as means ± SEM of four independent experiments, considering five microphotographs per experiment. Friedman test followed by One-way ANOVA followed by Bonferroni post-test: *p<0.05; **p<0.01 as depicted by lines.

### M(IL-4) cells are metabolically reprogrammed and become FM upon Mtb infection

As Mtb was shown to drive FM differentiation in both infected cells and uninfected bystander macrophages (2, 31, 32), we investigated whether the different activation profiles of macrophages differ in their ability to become FM this time upon Mtb infection. To this aim, M0, M(IFN-γ), M(IL-4) and M(IL-10) were infected (or not) with Mtb, and the accumulation of LBs was evaluated at 24 h post-infection. We confirmed that the infection with Mtb promoted LB accumulation at comparable levels in the different activation programs, including the M(IL-4) macrophages (Fig. 6A). Since we observed that STAT6 activation mediates the inhibition of FM formation in M(IL-4) macrophages, we next assessed whether Mtb infection resulted in a reduction in STAT6 phosphorylation in these cells. However, pSTAT6 was not reduced after Mtb infection of M(IL-4) cells (Fig. 6B), suggesting that Mtb targets another molecule downstream to STAT6 making M(IL-4) cells capable of accumulating LBs. Given the importance of the metabolic state in our model, we inferred that the M2-like metabolism was probably rewired due to infection. In the case of the glycolytic activity, we found that Mtb infection slightly increased glucose uptake and lactate release in both M0 and M(IL-4) cells (Fig. 6C-D). In addition, the expression levels of HIF-1α was higher in the Mtb-infected M(IL-4) cells compared to uninfected cells (Fig. 6E). As a proof-of-concept that M(IL-4) cells are metabolically re-programmed by Mtb infection, we assessed the accumulation of LBs in M(IL-4) macrophages in the presence of either a HIF-1α inhibitor (PX-478) or a FAO enhancer (L-carnitine). As expected, both treatments resulted in less accumulation of LBs triggered by Mtb infection in M(IL-4) macrophages (Fig. 6F-G and S7A). However, despite its effect on LB accumulation, L-carnitine per se was not enough to inhibit the bacterial growth compared to untreated M(IL-4) macrophages (Fig. 6H). Altogether, these findings demonstrate that the modulation of M(IL-4) metabolisms impacts on the acquisition of the foamy phenotype induced by Mtb.

**Figure 6.**
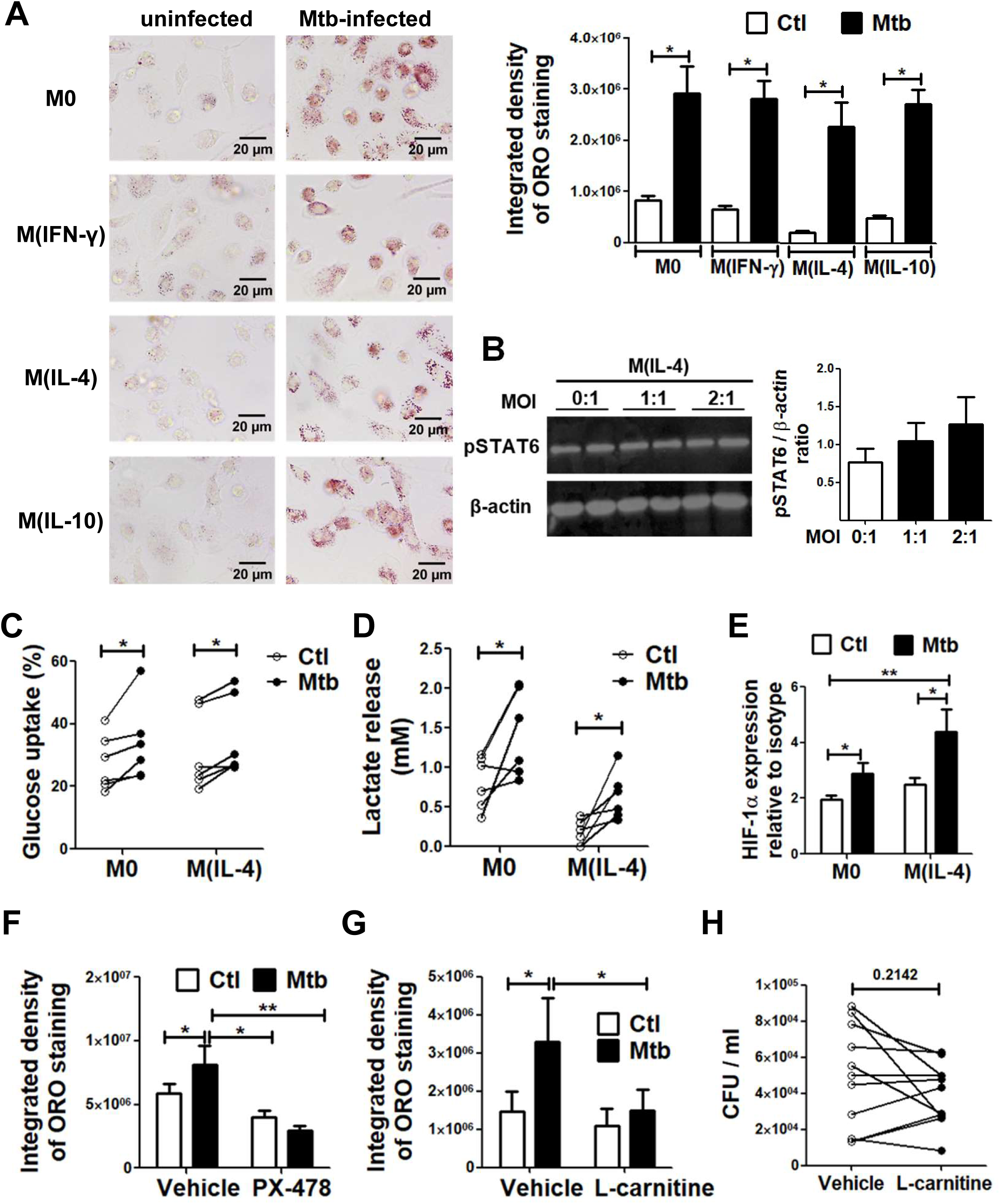
M(IL-4) cells are metabolically reprogrammed and become foamy upon Mtb infection. **(A)** ORO staining of M0, M(IFN-γ), M(IL-4) and M(IL-10) macrophages infected or not with Mtb. Representative images are shown in left panels (40× magnification) and the integrated density of ORO staining is shown in right panels. Values are expressed as means ± SEM of four independent experiments, considering five microphotographs per experiment. Wilcoxon Test: *p<0.05 for infected vs uninfected. **(B)** Analysis of pSTAT6 and β-actin protein expression level by Western Blot and quantification (n=4) in M(IL-4) cells infected or not with different multiplicity of Mtb infection. (C-F) M0 and M(IL-4) macrophages were infected or not with Mtb and the following parameters were measured: (C), glucose consumption (n=6), **(D)** lactate release (n=6) and, **(E)** HIF-1α expression (n=6). (F-G) ORO staining of M(IL-4) macrophages infected with Mtb and treated either with PX-478, a selective HIF-1α inhibitor **(F)** or with L-carnitine, a FAO enhancer (G). Values are expressed as means ± SEM of 6 independent experiments, considering five microphotographs per experiment. Friedman test followed by Dunn’s Multiple Comparison Test: *p<0.05; for experimental condition vs Ctl or as depicted by lines. **(H)** Intracellular colony forming units determined of M(IL-4) macrophages infected with Mtb and treated with L-carnitine at day 3 post infection.

## DISCUSSION

As a chronic inflammatory condition, TB entails the establishment of extensive metabolic reprogramming in both the host and the pathogen. One of the consequences of this metabolic adaptation is the formation of FM. Since FM have been associated with the bacilli persistence and tissue pathology (2, 31, 33, 34), we aimed to determine the impact of different activation/metabolic programs in human macrophages on the accumulation of LBs. By using the acellular fraction of TB-PE, known to induce FM formation in M0 and M(IL-10) macrophages (6), we demonstrated that STAT6 activation induced either by IL-4 or IL-13 prevents the accumulation of LBs in M(IL-4) macrophages. Importantly, we provide ex vivo evidence for the refractoriness of M(IL-4/IL-13) macrophages to become foamy by finding a negative correlation between the expression of CD209, a M(IL-4)-associated marker, and the numbers of LB-containing CD14^+^ cells isolated directly from the pleural cavity of TB patients, providing physiological relevance to our in vitro findings. Since tuberculous pleural effusions have, if any, very few bacilli content (35), we consider that the impact of potential in situ infection on the macrophage metabolic state, which may then lead M(IL-4) cells to become FM, might be very low. Additionally, while CD206 is also known to be induced by IL-4 and IL-13 (36), we did not find such an association between this marker and the numbers of FM. This can be due to the fact that its expression is also induced upon other activation programs such as M(IL-10), which are demonstrated to express high levels of CD206 (21, 37) and prone to synthetize and accumulate LBs (6). In fact, we have previously demonstrated that FM were formed after TB-PE treatment by increasing the biogenesis of LBs through ACAT up-regulation induced by the IL-10/STAT3 axis, generating cells accumulating LBs and displaying immunosuppressive properties (6). In this study, we observed that the STAT3 pathway is also induced in TB-PE-treated M(IL-4) macrophages (data not shown) resulting in ACAT induction, but as the newly formed LBs are rapidly disrupted in these macrophages, the foamy appearance was reduced drastically. In this regard, we showed that, while LBs are formed, they are quickly disrupted by an enhanced lipolytic activity induced in M(IL-4) macrophages, and those fatty acids produced through lipolysis are then incorporated into the mitochondria and used for FAO. Moreover, we demonstrated that cell population noted to rely on oxidative metabolism, such as AMs and M(IL-4) macrophages, are less prone to become FM. Hence, our findings are strengthened by previous results on the inhibition of the LBs accumulation in macrophages treated with resveratrol (38), since this phytoalexin is known to promote mitochondrial biogenesis and to increase the cellular respiratory capacity (39).

Interestingly, the inhibition of lipids oxidation in M(IL-4) macrophages results in diverting the fatty acid fate into triglycerides accumulation, leading M(IL-4) cells to form FM. Although the M2 program was noted to rely on FAO (13–16), the precise role of FAO in driving M2 polarization requires further study (40, 41). Herein we found that the exacerbated FAO activity prevents lipid accumulation and FM formation. Our results are in agreement with previous reports in the field of atherosclerosis, showing that enforcing CPT1a expression can reduce lipid accumulation (42), suggesting that the induction of FAO in foam cells could be of therapeutic potential. Also, in adipocytes, it has been demonstrated that IL-4 harbors pro-lipolysis capacity by inhibiting adipocyte differentiation and lipid accumulation, as well as by promoting lipolysis in mature cells to decrease lipid deposits (43).

Moreover, in agreement with our previous report and others, the differentiation of macrophages into FM (prior to infection) renders the cells more susceptible to the intracellular replication of Mtb, (6, 44, 45); in this study, we extended this notion to the M(IL-4) macrophage profile exposed to lipases inhibitors. We have previously shown that FM induced by TB-PE had immunosuppressive properties such as: i) high production of IL-10, ii) low production of TNF-α, iii) poor induction of IFN-γ producing T clones in response to mycobacterial antigens, and iv) more permissiveness to intracellular mycobacterial growth (6). Along with other data that associated FM with the persistence of infection (2, 32), these results support the idea that the generation of FM would have a negative impact for mycobacterial control; conversely, the refractoriness of macrophages to become foamy mediated by the IL-4/IL-13-STAT6 axis could have positive consequences for the host. Surprisingly, however, the infection with Mtb per se promoted the LB accumulation in M(IL-4) macrophages despite the activation of the IL-4/STAT6 axis. In fact, Mtb may induce the foamy phenotype by hijacking a metabolic pathway downstream to STAT6 activation. In addition, Mtb infection can decrease the lipolytic activity of macrophages at early time points (5, 34). Therefore, we propose that modulation of lipolysis is key for determining lipid accumulation in the context of Mtb infection. Of note, although we were able to inhibit FM differentiation upon Mtb infection by fostering the FAO pathway with L-carnitine, the cells were still susceptible to Mtb intracellular growth. Since macrophages having a more active FAO, and acquiring higher amounts of fatty acid, were reported as a preferred site for bacterial growth and survival (19), the effect of reducing the LB formation by L-carnitine may be counteracted by the exacerbation of the FAO metabolism, resulting in a net persistence of Mtb.

In this study, we demonstrated that M(IL-4) macrophages are reprogrammed metabolically by Mtb infection, leading to the formation of foam cells through the positive regulation of HIF-1α and/or the decrease in the FAO, without modulation in STAT6 phosphorylation. We consider that this mechanism constitutes a strategy of mycobacterial persistence. Indeed, a high lipid content guarantees the survival of the pathogen, and the activation of the IL-4 / STAT6 axis is associated with the establishment of a poorly microbicidal profile (9). According to our results, HIF-1α can be proposed as a molecular target to be modulated by Mtb infection impacting the LB accumulation. Previous reports have demonstrated that the infection with Mtb leads to glycolysis in bone-marrow derived macrophages (46, 47), in lungs of infected mice (48), and in lung granulomas from patients with active TB (49). Here, we showed that Mtb infection leads to HIF-1α activation and lactate release in M(IL-4) macrophages. Of note, Mtb infection induces the increase of HIF-1α expression in IFN-γ-activated macrophages (M1), which is essential for IFN-γ-dependent control of infection (50). Despite the fact that HIF-1α activation engages the microbicidal program in macrophages, we consider that Mtb can also take advantage of HIF-1α stabilization in M(IL-4) macrophages through the enhancement of LB accumulation, which is known to be detrimental for the inflammatory phenotype of macrophages (6, 51). It is also important to highlight that Mtb can reprogram the metabolic state of M(IL-4) macrophages without affecting STAT-6 activation. This means that the pro-inflammatory program, which could have been established upon HIF-1α activation, may also be tuned by the inhibitory signals driven by the IL-4/STAT-6 pathway. Hence, we speculate that foamy M(IL-4) macrophages can bear high bacillary loads despite HIF-1α activation, especially considering that LBs were described as a secure niche for mycobacteria conferring protection even in the presence of bactericidal mechanisms, such as respiratory burst (5). Moreover, our findings agree with a recent article by Zhang et al, in which they demonstrated that the deficiency of the E3 ligase von Hippel–Lindau protein (VHL), an enzyme that keeps HIF-1α at a low level via ubiquitination followed by proteasomal degradation, uplifted glycolytic metabolism in AM. Although not stressed by the authors, it is notable that these AM displayed a foamy phenotype unlike AM from WT mice (52), arguing that the uncontrolled activation of HIF-1α may contribute to FM formation. In line with this, it has recently been described that the activation of HIF-1α during late stages of infection with Mtb promotes the survival of infected FM (53), and that the activation of HIF-1α mediated by IFN-γ contributes to the formation of LBs in macrophages infected with Mtb (54). At least two hypotheses can explain how HIF-1α activation contributes to FM formation: 1) by suppressing FAO, and/or 2) by inducing endogenous fatty acid synthesis. Although in this study we did not address the endogenous synthesis of fatty acids of infected macrophages, we were able to demonstrate that both the inhibition of HIF-1α activity and the exacerbation of FAO partially decrease the acquisition of the foamy phenotype in M (IL-4) macrophages infected by Mtb. A possible link between the increase in HIF-1α and the decrease in FAO has been demonstrated in another experimental model, where both HIF-1α and HIF-2α mediate the accumulation of lipids in hepatocytes by reducing PGC-1α-mediated FAO (55).

Considering the collective impact of our findings, we postulate that the modulation of the metabolic pathways of macrophages, particularly those associated with lipid metabolism, could constitute a therapeutic target to improve the control of Mtb infection, complementing the current regimen of antibiotics (to make it briefer) and avoid the generation of bacterial multi-drug resistance.

## MATERIAL AND METHODS

### Chemical Reagents

DMSO, AS1517499, Etomoxir, L-carnitine, and Oil red O were obtained from Sigma-Aldrich (St. Louis, MO, USA); Lalistat was purchased from Bio-techne (Abingdon, UK), Tetrahydrolipstatin (Orlistat) from Parafarm (Buenos Aires, Argentina), PX-478 2HCL from Selleck Chemicals (Houston, USA) and DMOG from Santa Cruz, Biotechnology (Palo Alto, CA, USA).

### Bacterial strain and antigens

Mtb H37Rv strain was grown at 37°C in Middlebrook 7H9 medium supplemented with 10% albumin-dextrose-catalase (both from Becton Dickinson, New Jersey, United States) and 0.05% Tween-80 (Sigma-Aldrich). The Mtb γ-irradiated H37Rv strain (NR-49098) and its total lipidś preparation (NR-14837) were obtained from BEI Resource, USA. The RFP-expressing Mtb strain was gently provided by Dr. Fabiana Bigi (INTA, Castelar, Argentina).

### Preparation of human monocyte-derived macrophages

Buffy coats from healthy donors were prepared at Centro Regional de Hemoterapia Garrahan (Buenos Aires, Argentina) according to institutional guidelines (resolution number CEIANM-664/07). Informed consent was obtained from each donor before blood collection. Monocytes were purified by centrifugation on a discontinuous Percoll gradient (Amersham, Little Chalfont, United Kingdom) as previously described (6). After that, monocytes were allowed to adhere to 24-well plates at 5×10^5^ cells/well for 1 h at 37°C in warm RPMI-1640 medium (ThermoFisher Scientific, Waltham, MA). The medium was then supplemented to a final concentration of 10% Fetal Bovine Serum (FBS, Sigma-Aldrich) and human recombinant Macrophage Colony-Stimulating Factor (M-CSF, Peprotech, New Jork, USA) at 10 ng/ml. Cells were allowed to differentiate for 5-7 days. Thereafter, macrophage activation was induced by adding either IL-4 (20 ng/ml, Biolegend, San Diego, USA), IL-13 (20 ng/ml, Inmmunotools, Friesoythe, Germany), IL-10 (10 ng/ml, Peprotech), or IFN-γ (10 ng/ml, Peprotech) for further 48 h, resulting in M(IL-4), M(IL-13), M(IL-10) and M(IFN-γ) respectively.

### Preparation of pleural effusion pools

Pleural effusions (PE) were obtained by therapeutic thoracentesis by physicians at the Hospital F.J Muñiz (Buenos Aires, Argentina). The research was carried out in accordance with the Declaration of Helsinki (2013) of the World Medical Association and was approved by the Ethics Committees of the Hospital F.J Muñiz and the Academia Nacional de Medicina de Buenos Aires (protocol number: NIN-1671-12). Written informed consent was obtained before sample collection. Individual TB-PE samples were used for correlation analysis. A group of samples (n=10) were pooled and used for in vitro assays to treat macrophages. The diagnosis of TB pleurisy was based on a positive Ziehl-Neelsen stain or Lowestein–Jensen culture from PE and/or histopathology of pleural biopsy and was further confirmed by an Mtb-induced IFN-γ response and an adenosine deaminase-positive test (56). Exclusion criteria included a positive HIV test, and the presence of concurrent infectious diseases or non-infectious conditions (cancer, diabetes, or steroid therapy). None of the patients had multidrug-resistant TB. PE samples derived from patients with pleural transudates secondary to heart failure (HF-PE, n=5) were employed to prepare a second pool of PE, used as control of non-infectious inflammatory PE. The PE were collected in heparin tubes and centrifuged at 300 g for 10 min at room temperature without brake. The cell-free supernatant was transferred into new plastic tubes, further centrifuged at 12000 g for 10 min and aliquots were stored at −80°C. The pools were decomplemented at 56°C for 30 min and filtered by 0.22 µm in order to remove any remaining debris or residual bacteria.

### Foamy macrophage induction

Activated macrophages were treated with or without 20% v/v of PE, 10 μg/ml of Mtb lipids (BEI resources) or infected with Mtb (MOI 2:1) for 24 h. When indicated, cells were pre-incubated with either STAT6 inhibitor AS1517499 (100 nM, Sigma-Aldrich) for 30 min prior to IL-4 and TB-PE addition, orlistat (100 µM) and lalistat (10μM) for further 1 h after TB-PE incubation, etomoxir (3 and 10μM) for 30 min prior to TB-PE addition, DMOG (100μM) during TB-PE incubation and L-carnitine (1 and 10mM) for 48 h prior and during TB-PE addition. Foam cell formation was followed by Oil Red O (ORO) staining (Sigma-Aldrich) as previously described (6, 57) at 37°C for 1-5 min and washed with water 3 times. For the visualization of the lipid bodies, slides were prepared using the aqueous mounting medium Poly-Mount (Polysciences Inc, PA, -USA), observed via light microscope (Leica) and finally photographed using the Leica Application Suite software. For the determination of LBs size and numbers, images of ORO-stained cells were quantified with the ImageJ “analyze particles” function in thresholded images, with size (square pixel) settings from 0.1 to 100 and circularity from 0 to 1. For quantification, 10-20 cells of random fields (100x magnification) per donor and per condition were analyzed.

### Phenotypic characterization by flow cytometry

Macrophages were stained for 40 min at 4°C with fluorophore-conjugated mAb FITC-anti-CD36 (clone 5-271, Biolegend), or PE-anti-CD209 (clone 120507, R&D Systems, Minnesota, United States), and in parallel, with the corresponding isotype control antibody. After staining, the cells were washed with PBS 1X, centrifuged and analyzed by flow cytometry using a FACSCalibur cytometer (BD Biosciences, San Jose, CA, USA) The monocyte-macrophage population was gated according to its Forward Scatter and Size Scatter properties. The median fluorescence intensity (MFI) was analyzed using FCS Express V3 software (De Novo Software, Los Angeles, CA, USA).

### Western blots

Protein samples were subjected to 10% SDS-PAGE and the proteins were then electrophoretically transferred to Hybond-ECL nitrocellulose membranes (GE Healthcare, Amersham, UK). After blocking with 1% bovine serum albumin (BSA, DSF Lab, Córdoba, Argentina), the membrane was incubated with anti-human ACAT (1:200 dilution, SOAT; Santa Cruz) or anti-human pY641-STAT6 (1:200 dilution, Cell Signaling Technology, Danvers, MA, United States, clone D8S9Y) antibodies overnight at 4°C. After extensive washing, blots were incubated with HRP-conjugated goat anti-rabbit IgG (1:5000 dilution; Santa Cruz) antibodies for 1 h at room temperature. Immunoreactivity was detected using ECL Western Blotting Substrate (Pierce, ThermoFisher). Protein bands were visualized using Kodak Medical X-Ray General Purpose Film. For internal loading controls, membranes were stripped and then reprobed with anti-β-actin (1:2000 dilution; ThermoFisher, clone AC-15) or anti-STAT6 (1:1000 dilution; Cell Signaling Technology, clone D3H4) antibodies. Results from Western blot were analyzed by densitometric analysis (Image J software).

### Determination of metabolites

Glycerol release was determined following Garland and Randlès protocol (58) with some modifications. Briefly, cell-culture medium was removed and replaced by PBS containing 2% BSA fatty acid-free. Macrophages were incubated for 6 h at 37°C and supernatants were collected and assessed with the enzymatic kit TG Color GPO/PAP AA (Wiener, Argentina) according to the manufacturer’s instructions. In all cases, the absorbance was read using a Biochrom Asys UVM 340 Microplate Reader microplate reader and software.

### Determination of fatty acids uptake

Macrophages were incubated with 2.4 µM BODIPY 558/568 C12 (Red C12, Life Technologies, CA, USA) in PBS containing 0.1% BSA fatty acid-free, for 1 min at 37°C. Cells were immediately washed with cold PBS containing 0.2% BSA and run through the FACS machine using FL2 to determine fatty acids uptake.

### Measurement of cell respiration with Seahorse flux analyzer

Real-time oxygen consumption rate (OCR) in macrophages was determined with an XFp Extracellular Flux Analyzer (Seahorse Bioscience). The assay was performed in XF Assay Modified DMEM using 1.6×10^5^ cells/well and 3 wells per condition. Three consecutive measurements were performed under basal conditions and after the sequential addition of the following electron transport chain inhibitors: 3 μM oligomycin (OM), 1 μM carbonyl cyanide 4-(trifluoromethoxy)phenylhydrazone (FCCP), 0.5 μM rotenone (ROT) and 0.5 μM antimycin (AA). Basal respiration was calculated as the last measurement before addition of OM minus the non-mitochondrial respiration (minimum rate measurement after ROT/AA). Estimated ATP production designates the last measurement before addition of OM minus the minimum rate after OM. Maximal respiration rate (max) was defined as the OCR after addition of OM and FCCP. Spare respiration capacity (SRC) was defined as the difference between max and basal respiration.

### Quantitative RT-PCR

Total RNA was extracted with Trizol reagent (Sigma-Aldrich) and cDNA was reverse transcribed using the Moloney murine leukemia virus reverse transcriptase and random hexamer oligonucleotides for priming (Life Technologies). The expression of CPT1 was determined using PCR SYBR Green sequence detection system (Eurogentec, Seraing, Belgium) and the CFX Connect™ Real-Time PCR Detection System (Bio-Rad, CA, United States). Gene transcript numbers were standardized and adjusted relative to eukaryotic translation elongation factor 1 alpha 1 (EeF1A1) transcripts. Gene expression was quantified using the ΔΔCt method.

### Transmission Electron Microscopy (TEM)

M0 and M(IL-4) cells exposed to TB-PE were prepared for TEM analysis. For this purpose, cells were fixed in 2.5 % glutaraldehyde / 2 % paraformaldehyde (PFA, EMS, Delta-Microscopies) dissolved in 0.1 M Sorensen buffer (pH 7.2) during 2 h at room temperature and then they were preserved in 1 % PFA dissolved in Sorensen buffer. Adherent cells were treated for 1 h with 1% aqueous uranyl acetate then dehydrated in a graded ethanol series and embedded in Epon. Sections were cut on a Leica Ultracut microtome and ultrathin sections were mounted on 200 mesh onto Formvar carbon-coated copper grids. Finally, thin sections were stained with 1% uranyl acetate and lead citrate and examined with a transmission electron microscope (Jeol JEM-1400) at 80 kV. Images were acquired using a digital camera (Gatan Orius). For the determination of LBs size and number, TEM images were quantified with the ImageJ “analyze particles” plugins in thresholded images, with size (µm^2^) settings from 0.01 to 1 and circularity from 0.3 to 1. For quantification, 8-10 cells of random fields (1000x magnification) per condition were analyzed.

### Infection of human macrophages with Mtb

Infections were performed in the biosafety level 3 (BSL-3) laboratory at the Unidad Operativa Centro de Contención Biológica (UOCCB), ANLIS-MALBRAN (Buenos Aires), according to the biosafety institutional guidelines. Macrophages seeded on glass coverslips within a 24-well tissue culture plate (Costar) at a density of 5×10^5^ cells/ml were infected with Mtb H37Rv strain at a MOI of 2:1 during 1 h at 37°C. When indicated, PX-478 (100 µM) was added or not 1 h prior Mtb infection and renewed in fresh complete media after cell-washing. Then, extracellular bacteria were removed gently by washing with pre-warmed PBS, and cells were cultured in RPMI-1640 medium supplemented with 10 % FBS and gentamicin (50 µg/ml) for 24 h. In some experiments, L -carnitine (10 mM) was added or not after removing the extracellular bacteria. The glass coverslips were fixed with PFA 4% and stained with ORO, as was previously described.

### Measurement of bacterial intracellular growth in macrophages by CFU assay

Macrophages exposed (or not) to TB-PE, were infected with H37Rv Mtb strain at a MOI of 0.2 bacteria/cell in triplicates. After 4 h, extracellular bacteria were removed by gently washing four times with pre-warmed PBS. At different time points cells were lysed in 1% Triton X-100 in Middlebrook 7H9 broth. Serial dilutions of the lysates were plated in triplicate, onto 7H11-Oleic Albumin Dextrose Catalase (OADC, Becton Dickinson) agar medium for CFU scoring 21 days later.

### Visualization and quantification of Mtb infection

Macrophages seeded on glass coverslips within a 24-well tissue culture plate (Costar) at a density of 5 × 10^5^ cells/ml were infected with the red fluorescent protein (RFP) expressing Mtb CDC 1551 strain at a MOI of 5:1 during 2 h at 37°C. Then, extracellular bacteria were removed gently by washing with pre-warmed PBS, and cells were cultured in RPMI-1640 medium supplemented with 10% FBS for 48 h. The glass coverslips were fixed with PFA 4% and stained with BODIPY 493/503 (Life Technologies). Finally, slides were mounted and visualized with a FluoView FV1000 confocal microscope (Olympus, Tokyo, Japan) equipped with a Plapon 60X/NA1.42 objective, and then analyzed with the software ImageJ-Fiji. We measured the occupied area with RFP-Mtb (expressed as Raw Integrated Density) per cell in z-stacks from confocal laser scanning microscopy images. Individual cells were defined by BODIPY-stained cellular membranes which allow us to define the region of interests for quantification. For quantification 80-100 cells of random fields per donor per condition were analyzed.

### Mice

BALB/c male mice (8–12 weeks old) were used. Animals were bred and housed in accordance with the guidelines established by the Institutional Animal Care and Use Committee of Institute of Experimental Medicine (IMEX)-CONICET-ANM. All animal procedures were shaped to the principles set forth in the Guide for the Care and Use of Laboratory Animals (59).

### Culture of BMDMs and AMs

Femurs and tibia from mice were removed after euthanasia and the bones were flushed with RPMI-1640 medium by using syringes and 25-Gauge needles. The cellular suspension was centrifuged, and the red blood cells were removed by hemolysis with deionized water. The BMDMs were obtained by culturing the cells with RPMI-1640 medium containing L-glutamine, pyruvate, streptomycin, penicillin and β-mercaptoethanol (all from Sigma-Aldrich), 10% FBS and 20 ng/ml of murine recombinant M-CSF (Biolegend) at 37°C in a humidified incubator for 7–8 days. Differentiated BMDMs were re-plated on glass coverslips within 24-well tissue culture plates in complete medium and polarized or not with murine recombinant IL-4 (20 ng/ml, Miltenyi Biotec, CA, USA) for 48 h. AMs were harvested via bronchoalveolar lavage and plated on glass coverslips within a 24-well tissue culture plate in complete medium. Finally, BMDMs and AMs were stimulated with Mtb lipids (10 µg/ml) in the presence or not of DMOG (200μM) or etomoxir (3 µM) for 24 h and LBs accumulation was determined.

### Statistical analysis

Values are presented as means and SEM of a number of independent experiments or as indicated in figure legends. Independent experiments are defined as those performed with macrophages derived from monocytes isolated independently from different donors. For non-parametric paired data, comparisons were made using the Friedman test followed by Dunn’s Multiple Comparison Test or by two-tailed Wilcoxon Signed Rank depending on the numbers of experimental conditions to be compared. For parametric unpaired data, comparisons between two data sets were made by Mann Whitney test. For the analysis of the OCR measurements, t-test was applied; and for data obtained from cells derived from inbred mice, 1-way ANOVA followed by Bonferroni post-test was used. Correlation analyses were determined using the Spearman’s rank test. For all statistical comparisons, a p value <0.05 was considered significant.

## Acknowledgements

This work was supported by the Argentinean National Agency of Promotion of Science and Technology (PICT-2015-0055 to MCS and PICT-2017-1317 to LB), the Argentinean National Council of Scientific and Technical Investigations (CONICET, PIP 112-2013-0100202 to MCS), the Centre National de la Recherche Scientifique, and the Agence Nationale de Recherche sur le Sida et les Hépatites Virales (ANRS2014-CI-2, ANRS2014-049 and ANRS2018-1). The funders had no role in study design, data collection, and analysis, decision to publish, or preparation of the manuscript.

## Supplementary Figure Legends

**Supplementary figure 1. (A)** Median fluorescence intensity (MFI) of CD206, CD209, CD163, MerTK, CD86, and HLA-DR measured by flow cytometry on M(IL-4), M(IL-10) and M(IFN-γ) macrophages. Representative histograms and values of ten independent experiments are shown. Friedman test followed by Dunn’s Multiple Comparison Test: **p<0.01; ***p<0.001; as depicted by lines. **(B)** Analysis of pSTAT6, STAT6, pSTAT3, STAT3, pSTAT1, STAT1, and β-actin protein expression level by Western Blot and quantification in M(IL-4), M(IL-10) and M(IFN-γ) macrophages.

**Supplementary figure 2. (A)** MFI of CD209, CD206, and CD200R measured by flow cytometry on M0, M(IL-4), and macrophages polarized with different amounts of recombinant IL-13. Left panel represents the analysis of pSTAT6 and β-actin protein expression level M0, M(IL-4), and M(IL-13) by Western Blot (B-C) Representative images of ORO staining of macrophages polarized with either different doses of recombinant IL-4 **(B)** or IL-13 **(C)** for 48 h and exposed to TB-PE for further 24 h (40x magnification). **(D)** Cell viability of M(IL-4) macrophages exposed to different amounts of AS1517499 or vehicle. **(E)** Analysis of pSTAT6, STAT6, and β-actin protein expression level by Western Blot (right panel) and quantifications (left panels, n=4) in M0 and M(IL-4) macrophages exposed to AS1517499 or vehicle. Wilcoxon signed rank test: *p<0.05 as depicted by lines.

**Supplementary figure 3.** Correlation study between the MFI of CD206, HLA-DR, CD86, CD163, and MerTK cell-surface expression in CD14^+^ cells from TB pleural cavity and the percentage of lipid-laden CD14^+^ cells within the pleural fluids mononuclear cells (PFMC) (n=16) found in individual preparations of TB-PE. Spearman’s rank test.

**Supplementary figure 4. (A)** Glycerol release by M0, M(IL-4) and M(IL-13) macrophages (left panel, n=6). **(B)** Cell viability of M(IL-4) macrophages exposed to Orlistat (upper panel, n=4) and Lalistat (lower panel, n=4). **(C)** Glycerol release by M0 and M(IL-4) macrophages exposed or not to either Orlistat (left panel, n=6) or Lalistat (right panel, n=4). (D-E) Images of M(IL-4) macrophages infected with RFP-Mtb and stained with BODIPY 493/503 at 2 h and 48 h post-infection. **(D)** BODIPY intensity (left panel) and area occupied by RFP-Mtb (right panel) per cell in z-stacks from confocal laser scanning microscopy images at 2 h post-infection. Each determination represents individual cells of one donor. One way-ANOVA followed by Bonferroni test: ***p<0.001. **(E)** Area with RFP-Mtb per cell in z-stacks from confocal laser scanning microscopy images. Values are expressed as means of 80-100 cells in four independent experiments. Friedman followed by Dunn’s Multiple Comparison Test: *p<0.05 as depicted by lines.

**Supplementary figure 5. (A)** Cell viability of M(IL-4) macrophages exposed to different amounts of Etomoxir (n=4). **(B)** Cell viability of M0 macrophages exposed to different amounts of L-carnitine (n=8). **(C)** MFI of CD209, and CD200R measured by flow cytometry on M(IL-4) treated or not with TB-PE. **(D)** Analysis of pSTAT6, STAT6, and β-actin protein expression level by Western Blot (left panel) in M(IFN-γ), M(IL-4), and M(IL-10) macrophages treated or not with TB-PE. Quantifications in M(IL-4) cells (right panels, n=4). Wilcoxon signed rank test: *p<0.05. **(E)** CFSE-labelled apoptotic neutrophils (PMN) were cocultured with M(IL-4) macrophages treated or not with TB-PE for 1 h, washed stringently to remove free PMN, and the percentage of macrophages positive for CFSE was assessed. Representative dot blots showing CFSE labeling of apoptotic PMN, M(IL-4), and cocultures are shown. Blue gates represent mainly CFSE labeling of apoptotic PMN while red gates comprise mainly macrophages. Lower panels: Annexin V positive vs propidium iodide negative of apoptotic PMN (left panel) and percentages of M(IL-4) treated or not with TB-PE macrophages ingesting apoptotic PMN (right panel). M(IFN-γ) macrophages were also tested for comparison.

**Supplementary figure 6. (A)** Human macrophages were left untreated (M0) or polarized with IL-4 (M(IL-4)) for 48h, treated or not with the acellular fraction of TB pleural effusions (TB-PE) in the presence or not of DMOG (100 µM) for 24 h and then stained with Oil Red O (ORO). Representative images are shown (40× magnification). **(B)** Cell viability of M0 macrophages exposed to different amounts of DMOG. **(C)** Left panel: Representative images (40× magnification) of ORO staining of murine AM treated or not with Mtb lipids in the presence of either DMOG (200 µM) or Etomoxir (3 µM). Right panel: Lactate release and glucose consumption by AM treated or not with Mtb lipids in the presence of DMOG for 24 h (n=3). One-way ANOVA followed by Bonferroni’s Multiple Comparison Test: *p<0.05; **p<0.01, as depicted by lines. **(D)** Representative images (40× magnification) of ORO staining of murine bone marrow derived macrophages murine (BMDM) unpolarized or polarized towards M(IL-4), exposed or not to Mtb lipids.

**Supplementary figure 7.** Lactate release by M(IL-4) macrophages infected or not with Mtb and treated with PX-478 or its vehicle (DMSO) for 24 h (n=4). Friedman followed by Dunn’s Multiple Comparison Test: *p<0.05 as depicted by lines.

